# MIR2052HG regulates ERα levels and aromatase inhibitor resistance through LMTK3 by recruiting EGR1

**DOI:** 10.1101/513879

**Authors:** Junmei Cairns, James N. Ingle, Krishna R. Kalari, Lois E. Shepherd, Michiaki Kubo, Matthew P. Goetz, Richard M. Weinshilboum, Liewei Wang

## Abstract

Our previous GWAS using the MA.27 aromatase inhibitors (AIs) adjuvant trial identified SNPs in the lncRNA *MIR2052HG* associated with breast cancer free interval. Here we report that MIR2052HG depletion in breast cancer cells results in a decrease in LMTK3 expression and cell growth. Mechanistically, MIR2052HG interacts with EGR1 and facilitates its recruitment to the LMTK3 promoter. LMTK3 sustains ERα levels by reducing PKC activity, resulting in increased ESR1 transcription mediated through AKT/FOXO3 and reduced ERα degradation mediated by the PKC/MEK/ERK/RSK1 pathway. MIR2052HG regulated LMTK3 in a SNP- and aromatase inhibitor - dependent fashion: the variant SNP increased EGR1 binding to LMTK3 promoter in response to androstenedione, relative to wild-type genotype, a pattern that can be reversed by aromatase inhibitor treatment. Finally, LMTK3 overexpression abolished the effect of MIR2052HG on PKC activity and ERα levels. These results reveal a direct role of MIR2052HG in LMTK3 regulation and raise the possibility of targeting MIR2052HG or LMTK3 in ERα-positive breast cancer.

## Introduction

Estrogens have long been recognized to be important for stimulating the growth of estrogen receptor α (ERα) positive breast cancer, a subtype that represents a large proportion of breast cancer patients. Estrogen action is mediated by ERα. Approximately 70% of breast cancers are ERα positive and rely on estrogen signaling to stimulate their growth and survival [1,2]. Its presence in breast tumors is routinely used to predict response to endocrine therapies that target ERα, estrogen production or estrogen signaling. AIs suppress estrogen synthesis in postmenopausal women by targeting the aromatase enzyme, which converts precursor hormones to estrogens. The third-generation AIs (i.e., exemestane, anastrozole, and letrozole) have largely replaced tamoxifen as the preferred treatment for ERα positive breast cancer in postmenopausal women with early stage breast cancer because of their superior efficacy over tamoxifen [3,4]. However, both de novo and acquired resistance to AIs can occur, resulting in relapse and disease progression. It is estimated that approximately 30% of ER-positive breast cancer receiving adjuvant AI treatment eventually develop resistance [5-7], while nearly all patients develop resistance in the metastatic setting. The mechanisms for endocrine therapy resistance are complex and one mechanism includes dysregulation of ERα expression, encoded by *ESR1* [8].

ERα is a member of the nuclear receptor superfamily of ligand-activated transcription factors [9], which regulates gene expression through direct binding to estrogen response elements (EREs) in promoters of estrogen-regulated genes and indirectly through recruitment to gene promoters by interaction with other transcription factors [10]. Previous studies have reported that ESR1 is up-regulated during estrogen deprivation adaptation [11]. Overproduction of ERα leads to an enhanced response to low concentrations of estrogen, which is responsible for the acquisition of AI resistance or postmenopausal tumorigenesis [12,13]. In these AI-resistant tumors, ERα is either hypersensitive to low levels of estrogens [14] activated in a ligand-independent manner by phosphorylation via kinases in the growth factor receptor signaling pathways, or by acquired somatic *ESR1* mutations [15,16]. ERα phosphorylation aids in regulating the transcriptional activity and turnover of ER by proteasomal degradation. Of particular importance are Ser118 and Ser167, which locate within the activation function 1 region of the N-terminal domain of ERα and are regulated by multiple signaling pathways [17-20]. The phosphorylation at Ser118 can be mediated by mitogen-activated protein kinase (MAPK) activation and induces ERα activity [15,21], whereas Ser167 can be phosphorylated by p90RSK [22,23] and plays a role in Lemur Tyrosine Kinase 3 (LMTK3)-mediated ERα stabilization [24,25]. LMTK3 has been implicated in both de *novo* and acquired endocrine resistance in breast cancer [26]. The phosphorylation of ERα at S167 is positively associated with pMAPK and pp90RSK in breast cancer patients and a predictor of better prognosis in primary breast cancer with reduced relapse and better overall survival [27].

Our previous genome-wide association study (GWAS) used samples from the Canadian Cancer Trials Group MA.27, the largest AI breast cancer adjuvant endocrine therapy trial (4,406 controls without recurrence of breast cancer and 252 cases with recurrence). In that study, we identified common SNPs in a long noncoding (lnc) RNA, MIR2052HG, that were associated with breast cancer free interval (HR= 0.37, P= 2.15E-07) [28]. The variant SNPs (minor allele frequency [MAF]= 0.32 to 0.42) were associated with lower MIR2052HG and ERα expression in the presence of AIs, and two of the top SNPs, rs4476990 and rs3802201, were located in or near an ERE [28]. MIR2052HG appeared to affect ERα expression both by promoting AKT/FOXO3-mediated ESR1 transcription regulation and by limiting ubiquitin-mediated ERα degradation [28]. However, the underlying mechanisms by which MIR2052HG regulates ESR1 transcription and ERα degradation remain unknown.

LncRNAs are transcripts with no protein-coding functions. Accumulating evidence suggests that lncRNAs play critical roles in regulating a wide range of cellular processes through affecting various aspects of protein, DNA, and RNA expression and interactions [29-31]. Several lncRNAs have been implicated in breast cancer. For example, UCA1 is an oncogene in breast cancer that can induce tamoxifen resistance [32]. LncRNA HOTAIR is positively correlated with tamoxifen resistance [33]. In the current study, we sought to further investigate the mechanism of MIR2052HG action in the regulation of ERα and AI resistance. We found that MIR2052HG directly interacts with the early growth response protein 1 (EGR1) protein to enhance LMTK3 transcription and thus sustained ESR1 expression and stabilized ERα protein.

## Results

### MIR2052HG regulates LMTK3 expression

We previously reported that MIR2052HG sustained ERα levels by promoting AKT/FOXO3 mediated upregulation of ESR1 transcript and by limiting proteasome-dependent degradation of ERα protein [28]. However, the mechanism involved in the regulation of MIR2052HG-mediated AKT activation and ERα ubiquitination remains unknown. Kinome screening previously identified LMTK3 as a potent ERα regulator, acting by decreasing the activity of protein kinase C (PKC) and the phosphorylation of AKT (Ser473), resulting in increased binding of FOXO3 to the *ESR1* promoter [24]. LMTK3 also protected ERα from proteasome mediated degradation [24]. Given that the effects of LMTK3 on ERα were similar to our observations with MIR2052HG [28], we hypothesized that MIR2052HG might regulate LMTK3 to mediate ERα levels and in turn, response to AIs.

Previous studies have demonstrated that lncRNAs can function in *trans* to regulate expression of protein-coding genes; therefore, we examined the possibility that MIR2052HG may facilitate AI resistance by regulating LMTK3 expression. Consistent with this hypothesis, we found that knockdown of MIR2052HG using a pooled antisense oligonucleotides (ASO) in human ER positive CAMA-1 breast cancer cells resulted in a dramatic decrease of LMTK3 expression (Fig 1A). A similar effect was also observed in an aromatase overexpressing cell line, MCF7/AC1 [34] (Fig 1B). We also observed that the changes in mRNA levels were confirmed at the protein level by the western blot analysis (Fig 1C), supporting the notion that MIR2052HG regulates LMTK3 expression. To determine whether LMTK3 is a major downstream target of MIR2052HG in regulating AI response, we first determined the transcriptome changes in MIR2052HG -knockdown cells and collected published RNA-seq data after LMTK3-knockdown [26]. Analysis of the RNA-seq data indicated that the changes induced by MIR2052HG knockdown and LMTK3 knockdown showed a large number genes overlapped, especially almost 1/3 genes regulated by LMTK3 were also regulated by MIR2052 (Fig 1D, Table EV1). The common dysregulated genes in both knockdowns included cell cycle genes, oocyte maturation and oocyte meiosis genes (Table EV1).

**Figure 1.**
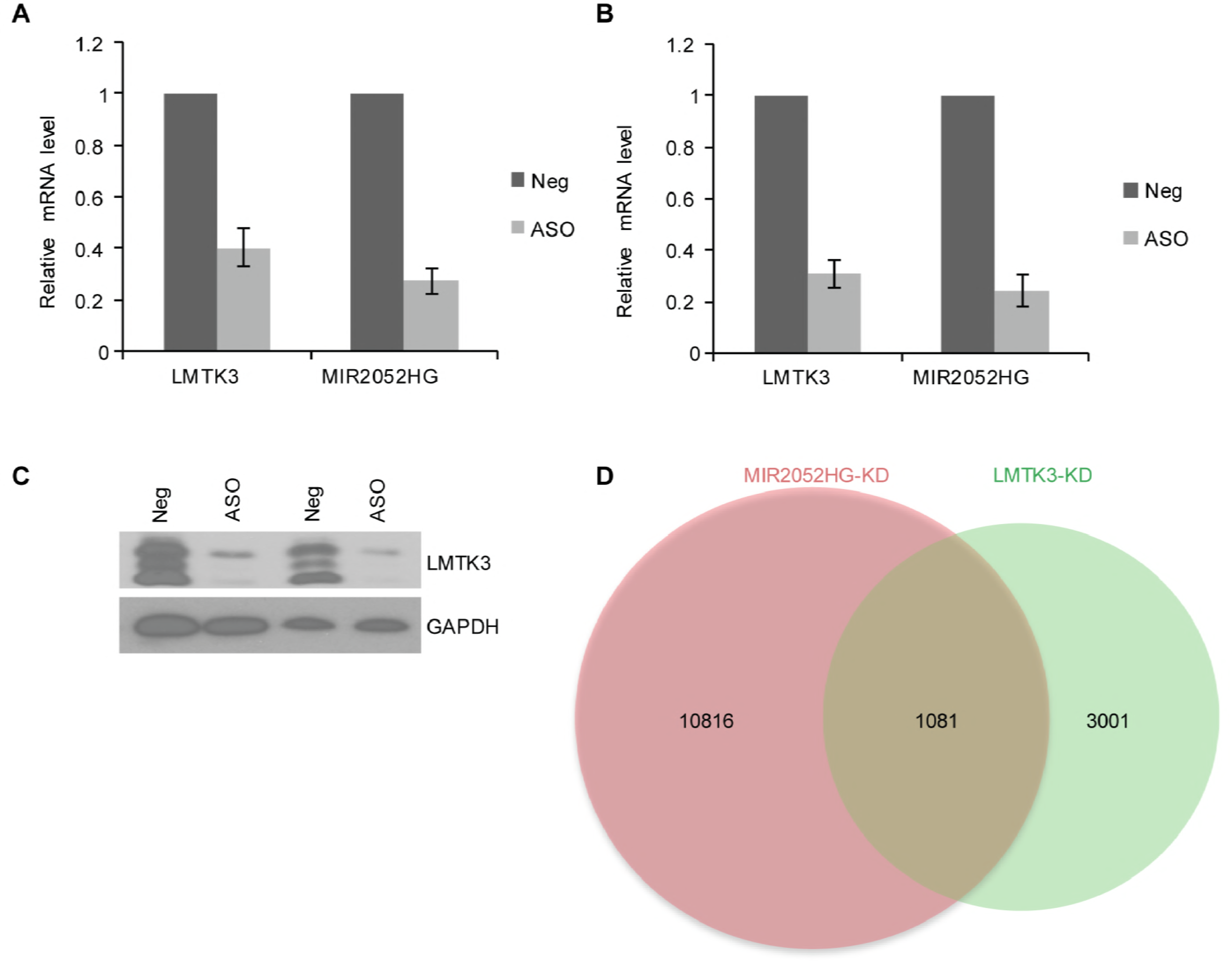
LMTK3 mRNA and protein expression in MIR2052HG knockdown cells. A, B Relative mRNA expression of LMTK3 after knockdown of MIR2052HG using pooled antisense oligonucleotides (ASO) in MCF7/AC1 (a) and CAMA-1 (b) cells. Error bars represent SEM; ** P< 0.01 compared to baseline (negative control). C Western blots analysis of LMTK3 after knocking down MIR2052HG in MCF7/AC1 and CAMA-1 cell lines. D Venn diagram shows that genes affected by MIR2052HG knockdown largely overlap with those affected by LMTK3 knockdown.

To further define the relationship between MIR2052HG and LMTK3, we transfected LMTK3 overexpressing constructs into ERα positive breast cancer cells with MIR2052HG -knockdown, followed by cell growth and colony forming assays. The cell proliferation and colony formation analysis demonstrated that the cell growth defect caused by downregulation of MIR2052HG could be successfully rescued by LMTK3 overexpression (Fig 2A-D), indicating that LMTK3 is a major target that mediates the MIR2052HG regulation on cell growth in ER positive breast cancer.

**Figure 2.**
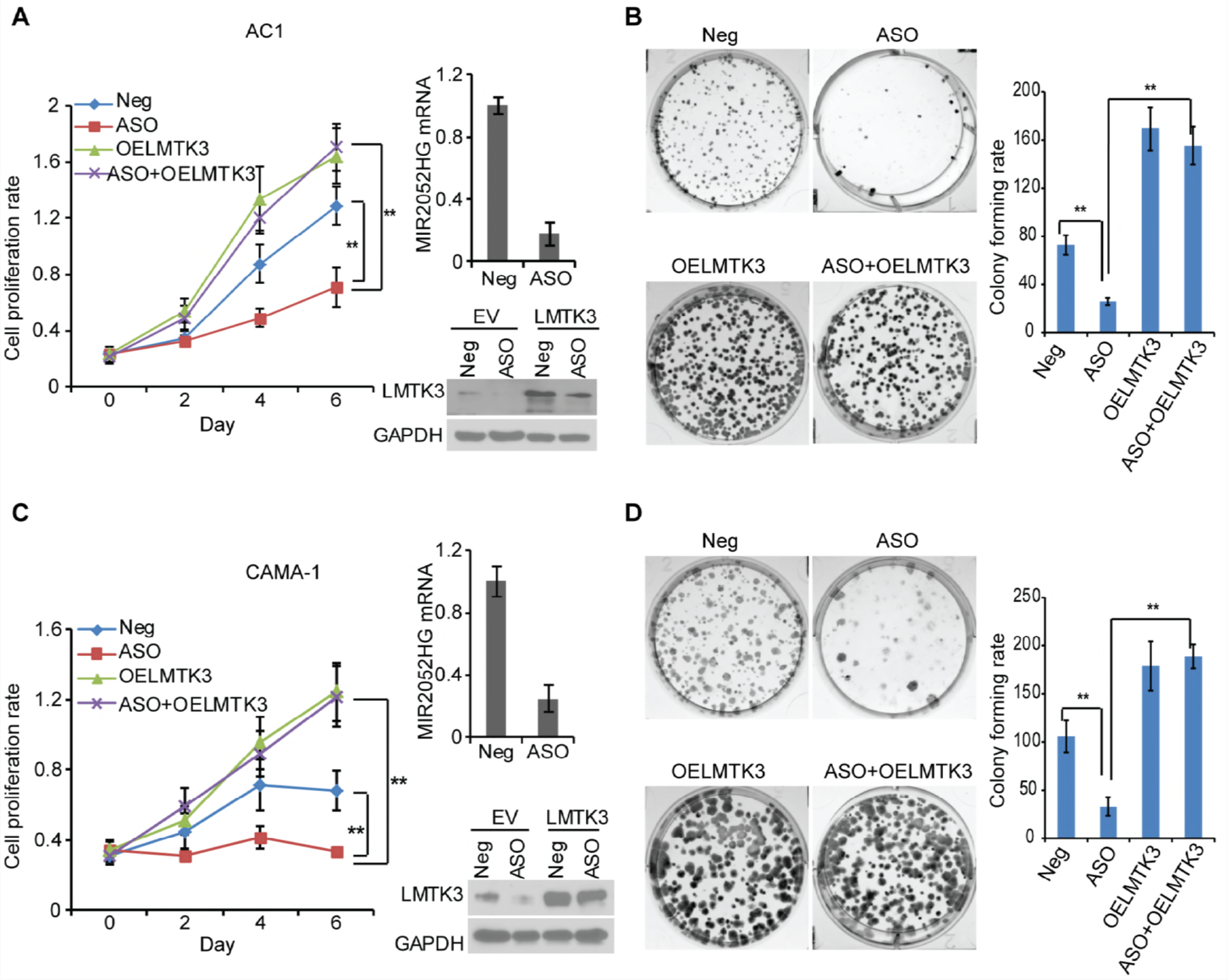
MIR2052HG regulates breast cancer cell growth through LMTK3-mediated signaling. A-D Overexpression of LMTK3 in MIR2052HG knocked-down MCF7/AC1 (A, B) and CAMA-1 (C, D) cells reversed the phenotypes of cell proliferation (A, C) and colony formation (B, D). Knockdown efficiency was determined by qRT-PCR. Overexpression of LMTK3 was determined by western blotting (A, C; right panel). Representative pictures of colony formation from three independent experiments are shown (B, D; left panel). The colony formation rates are quantified (B, D; right panel). Error bars represent SEM; ** p< 0.01.

### LMTK3 mediates MIR2052HG -regulation of ESR1 transcription and ERα protein stability

Previous studies demonstrated that MIR2052HG regulates ERα expression through transcription regulation of *ESR1* and ER protein degradation [24,28]. However, the direct target of MIR2052HG in ERα regulation has not been fully elucidated. Therefore, we examined the role of LMTK3 in ER**α** regulation. Our previous report indicated that the effect of MIR2052HG on ESR1 transcript is through AKT/FOXO3 [24]. In MCF7/AC1 and CAMA-1 cells, down-regulation of MIR2052HG reduced *ESR1* mRNA levels by promoting AKT-mediated down-regulation of FOXO3 protein level, a transcription factor known to be involved in ESR1 transcription (Fig 3A, B). Whereas LMTK3 overexpression rescued the downregulation of ERα mRNA induced by MIR2052HG silencing (Fig 3A, B). LMTK3 overexpression resulted in a decrease in phosphorylated AKT (at Ser473), and an increase in FOXO3protein level but not mRNA level (Fig 3A, B, Fig EV1A). At the protein level, ERα protein was reduced by MIR2052HG knockdown, whereas LMTK3 overexpression stabilized ERα (Fig 3A, B, right panel). Our previous results showed that MIR2052HG also regulated ERα stability by regulating its proteasome dependent degradation process. Here, LMTK3 overexpression could increase protein level by decreasing ERα ubiquitination (Fig 3C). Together, these data indicate that LMTK3, downstream of MIR2025HG, mediated MIR2052HG effect on the regulation of ESR1 transcription and ER**α** protein stability. Using the TCGA data set, LMTK3 showed higher expression level in ER positive breast cancer patients compared with other subtypes (Fig 3D), and RNA expression levels were also independently associated with disease-free survival and overall survival (Fig 3E).

**Figure 3.**
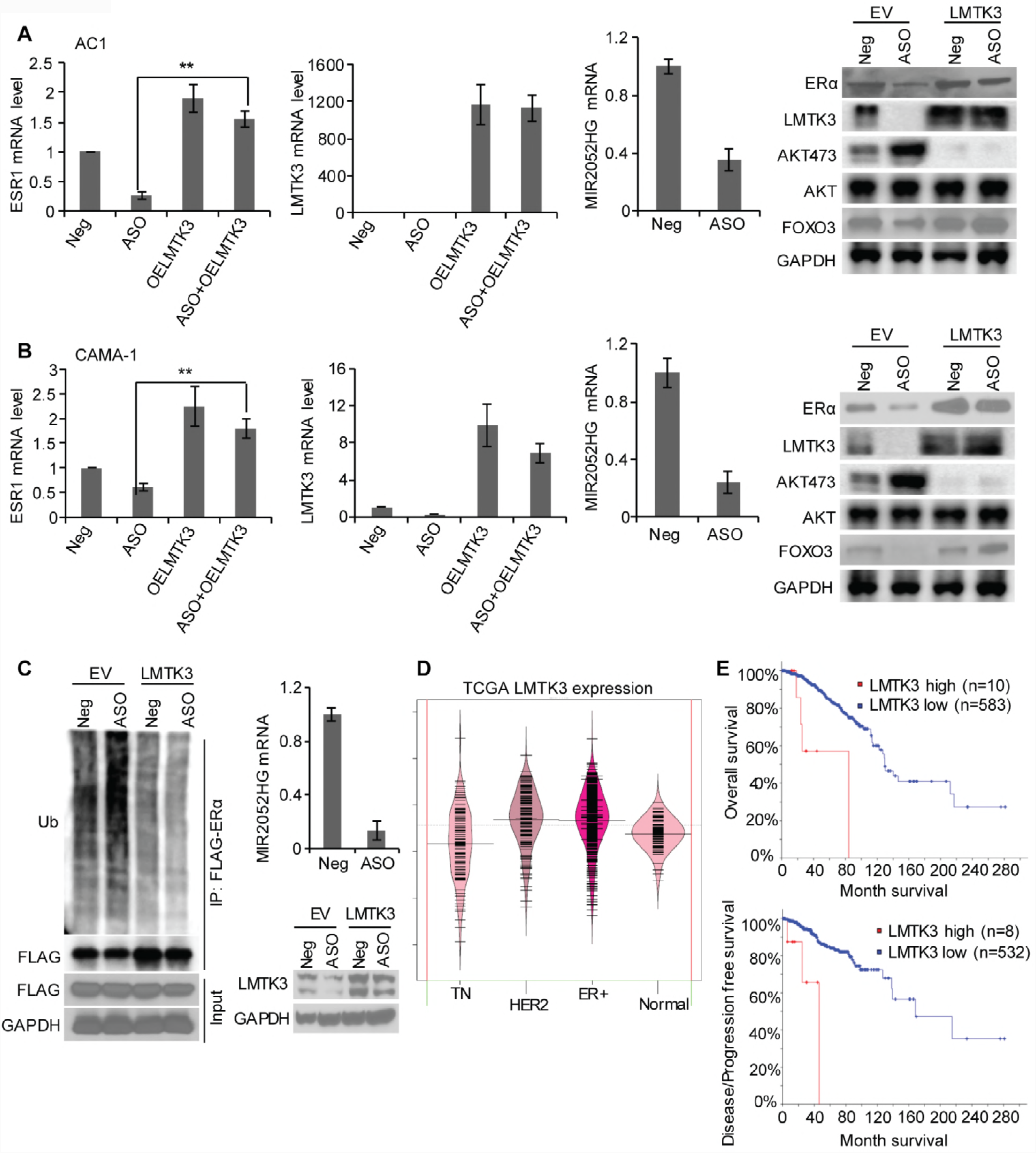
LMTK3 mediated MIR2052HG effect on regulation of ESR1 transcription and ERα protein stability. A, B Overexpression of LMTK3 in MIR2052HG knocked-down MCF7/AC1 (A) and CAMA-1 (B) cells reversed ERα protein and mRNA levels, and in turn, decreased AKT phosphorylation and FOXO3 level. Knock down efficiency was determined by qRT-PCR. Overexpression of LMTK3 was determined by western blot analysis. C Overexpression of LMTK3 in MIR2052HG knocked-down cells reduced the ubiquitination of ERα. 293T cells were transfected with HA-Ub plasmid and FLAG-ERα plasmid, and then transfected with either the MIR2052HG ASOs or LMTK3 plasmid followed by MG132. ERα proteins were immunoprecipitated and analyzed by western blot analysis (left panel). Knock down efficiency was determined by qRT-PCR. Overexpression of LMTK3 was determined by western blot (right panel). Error bars represent SEM; ** p< 0.01. D LMTK3 expression in TCGA breast cancer patients. E Kaplan-Meier plots demonstrated the associations between LMTK3 expression level and overall survival (P = 3.927e-5) as well as disease-free survival (P = 9.587e-5).

### MIR2052HG regulates ERα protein degradation through the LMTK3/PKC/MEK/ERK/RSK1 pathway

Next, we investigated the mechanisms involved in MIR2052HG and LMTK3 regulation of ERα protein degradation. ERα phosphorylation, especially increased phosphorylation of ERα at Ser167, has been implicated in ERα proteasome-mediated degradation [24]. To explore the mechanism, we first determined the level of ERα pSer167 in MIR2052HG knockdown ERα positive breast cancer cells. ERα pSer167 levels increased with MIR2052HG knockdown, despite decreased total ERα amounts (Fig 4A). Furthermore, we observed that knocking down MIR2052HG increased wild type (WT) ERα ubiquitination, but not the mutant ERα with serine 167 to alanine (S167A) (Fig 4B), confirming the involvement of ERα Ser167 in ubiquitin-dependent and proteasome-mediated degradation. The phosphorylation of ERα at Ser167 is regulated by pp90 (RSK1) [22], which is activated by MAPK [35]. We thus hypothesized that MEK/ERK/p90RSK1 might be the signaling pathway that mediates ERα phosphorylation at Ser167 upon MIR2052HG knockdown. We first tested the effect of knockdown of MIR2052HG on MEK/ERK/p90RSK1 activity. As shown in Figure 4A, in MCF7/AC1 and CAMA-1 cells transfected with MIR2052HG ASO, coinciding with increased ERα pSer167, pMEK, pERK and pRSK1levels were also increased, indicating an increased MEK/ERK/p90RSK1 activity. We then examined the role of LMTK3 in MIR2052HG-mediated regulation of MEK/ERK/p90RSK1 activity, and found that LMTK3 overexpression abolished increased pMEK/pERK/p90RSK1 levels caused by MIR2052HG silencing (Fig 4C, D). These data further imply that MIR2052HG regulates LMTK3 expression, which then influences MEK/ERK/p90RSK1 activity, regulating ERα protein levels.

**Figure 4.**
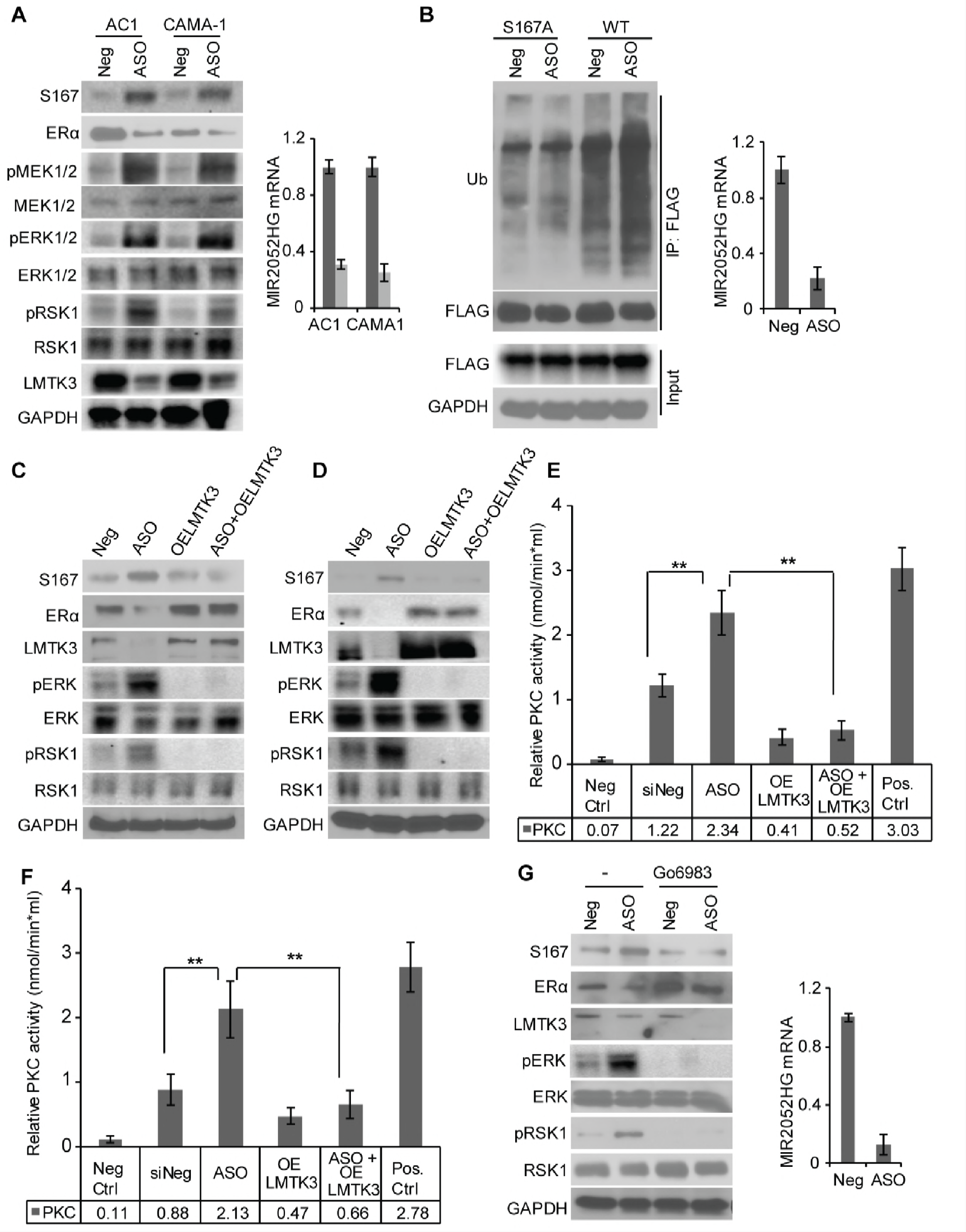
MIR2052HG regulates ERα protein stability through MEK/ERK/RSK1 pathway. A Knockdown of MIR2052HG increased phosphorylation of MEK, ERK, RSK1, as well as ERα S167 and decreased LMTK3 total level in MCF7/AC1 and CAMA-1 cells. B Knockdown of MIR2052HG promoted the ubiquitination of wild type ERα, but not ERα S167A mutant. 293T cells were transfected with HA-Ub plasmid and FLAG-ERα or FLAG-ERα S167A plasmid, and then transfected with either the MIR2052HG specific ASOs or the negative control ASO followed by MG132. Wild type or S167A mutant ERα proteins were immunoprecipitated and analyzed by western blot. Knock down efficiency in 293T cells was determined by qRT-PCR. C, D Overexpression of LMTK3 in MIR2052HG knocked-down MCF7/AC1(C) and CAMA-1 (D) cells reversed ERα protein levels and the phosphorylation of MEK, ERK, RSK1, and ERα S167. Overexpression of LMTK3 was determined by western blot analysis. E, F PKC kinase assays examining the effect of MIR2052HG and LMTK3 on the catalytic activity of PKC in MCF7/AC1 (E) and CAMA-1 (F) cells. Error bars represent SEM of two independent experiments in triplicate; ** p< 0.01. G Effects of MIR2052HG silencing on ERα protein levels in the presence of a PKC inhibitor (Go 6983).

As protein kinase C (PKC) has been implicated to play a role in ERα protein degradation [36] and AKT –FOXO3 regulation [37], and LMTK3 inhibits PKC activity [24], we examined the effects of MIR2052HG and LMTK3 on PKC. *In vitro* kinase assays indicated that downregulation of MIR2052HG increased PKC activity (Fig 4E, F, Fig EV1B), whereas overexpression of LMTK3 decreased PKC activity (Fig 4E, F, Fig EV1B). In addition, LMTK3 overexpression dramatically reduced PKC activity that was induced by MIR2052HG silencing (Fig 4E, F, and Fig EV1B). Inhibition of PKC with the Go 6983 inhibitor reduced MEK/ERK/p90RSK1 activity and ERα pSer167, which in turn, partially rescued ERα levels (Fig 4G), suggesting that MIR2052HG regulated ERα protein level through the axis of LMTK3/PKC/MEK/ERK/RSK1. Our findings also confirmed that MIR2052HG effects on AKT/ FOXO3 activation and downstream ESR1 mRNA level were through the regulation of LMTK3/PKC pathway (Figs 3-4).

### MIR2052HG contributes to LMTK3 transcription by facilitating EGR1 recruitment

Next, we wanted to address how MIR2052HG regulates LMTK3 transcription. First, we examined the localization of the MIR2052HG RNA transcript. RNA fluorescent in situ hybridization (FISH) demonstrated that MIR2052HG localized to a limited number of nuclear foci (one to two spots in most cases), suggesting that MIR2052HG had limited targets. We also checked the genomic location of the MIR2052HG transcript by RNA-DNA dual FISH, and the results showed that the MIR2052HG transcript was located at the LMTK3 gene locus (Fig 5A, Fig EV2). Taken together, these data suggest that MIR2052HG is likely involved in LMTK3 transcription.

**Figure 5.**
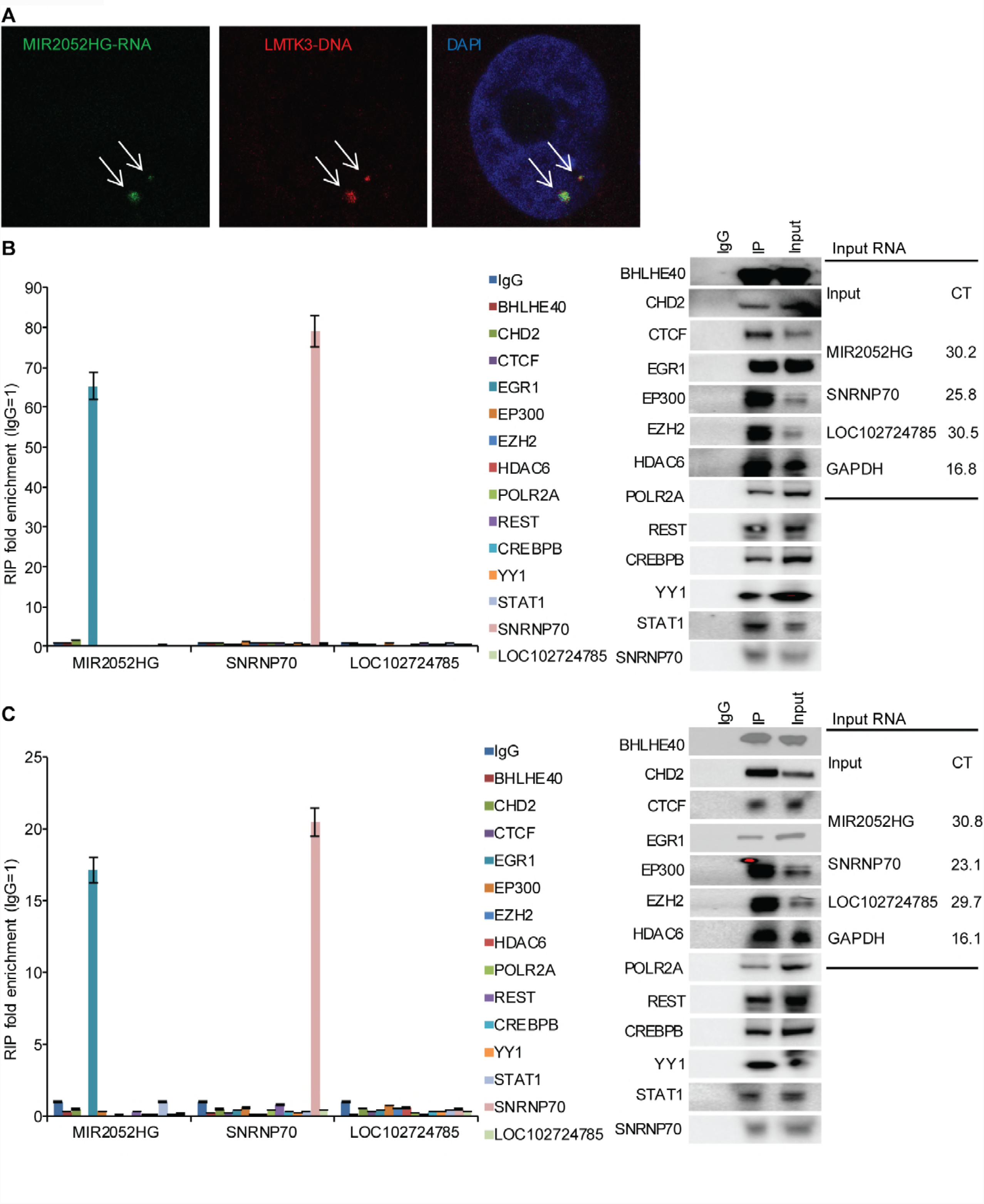
MIR2052HG regulates LMTK3 transcription by facilitating EGR1 recruitment to the LMTK3 promoter. A Dual RNA-DNA-FISH demonstrates that MIR2052HG transcripts (green signal) are localized onto the LMTK3 gene locus (red signal). B, C EGR1 antibody immunoprecipitates MIR2052HG, but not lncRNA LOC102724785 in AC1 (B) and CAMA-1 (C) cells. Error bars represent SEM of two independent experiments in triplicate; ** p< 0.01.

LMTK3 expression could be activated by several transcription factors based on the ENCODE database, including BHLHE40, CHD2, CTCF, EGR1, EP300, EZH2, HDAC6, POLR2A, REST, CREBBP, YY1, and STAT1. Therefore, we asked whether any of these transcription factors, together with MIR2052HG might be involved in the regulation of MIR2052HG expression. Immunoprecipitation followed by qRT-PCR analysis demonstrated that MIR2052HG was significantly enriched in the EGR1 immunoprecipitates (Fig 5B, C). The enrichment of MIR2052HG by the EGR1 antibody was specific, as the antibody did not pull down another lncRNA, LOC102724785 (Fig 5B, C). These data suggest that MIR2052HG regulation of LMTK3 transcription involves EGR1.

EGR1 was highly expressed in the Cancer Genome Atlas (TCGA) [38] ER positive breast cancer patients (Fig EV3). We then confirmed EGR1 regulation of LMTK3 gene expression in the MCF7/AC1 and CAMA-1 cells. Knockdown of EGR1 reduced LMTK3 mRNA level (Fig 6A, B). To examine whether binding of EGR1 to the *LMTK3* promoter requires MIR2052HG, we first mapped the binding locations of EGR1 on the *LMTK3* gene locus (Fig 6C, chr19:48994366-48994811, chr19:48996320-48996559, chr19:49015095-49015334). Chromatin immunoprecipitation (ChIP) assays demonstrated that EGR1 bound to the promoter region of *LMTK3* (Fig 6D, E). Importantly, knocking down MIR2052HG reduced the EGR1binding to the *LMTK3* promoter region (Fig 6D, E) without significant effect on the binding of EGR1 to other EGR1 targets (Figs EV4 and EV5). Furthermore, MIR2052HG failed to locate in the *LMTK3* gene locus in EGR1 knockdown cells (Fig 6F).

**Figure 6.**
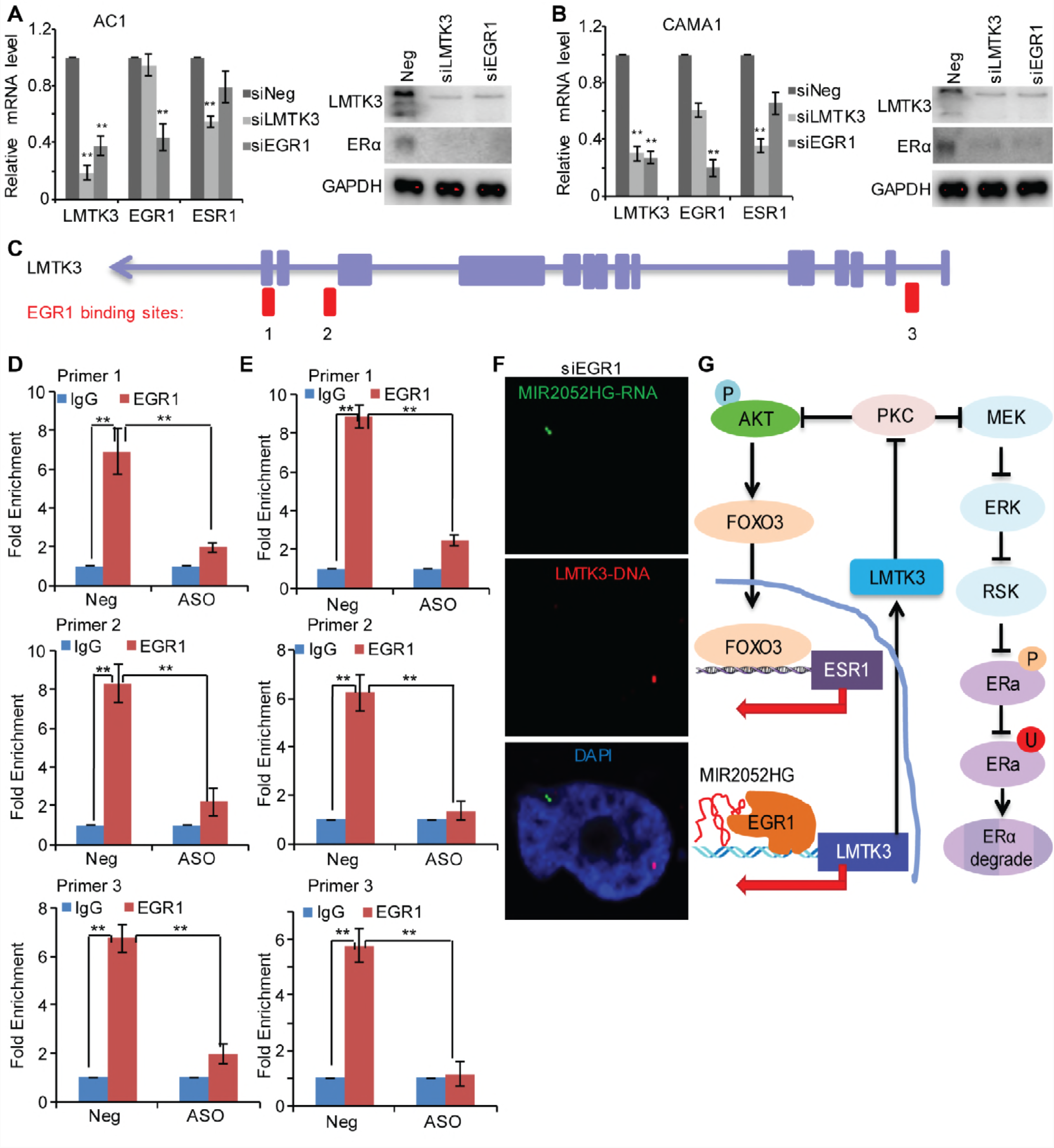
MIR2052HG regulates LMTK3 transcription by regulating EGR1 binding to its motif in LMTK3 gene. A, B EGR1 regulates LMTK3 expression in AC1 (A) and CAMA-1 (B) cells. C The EGR1 binding sites at the genomic location of the LMTK3 gene locus are indicated in the diagram. D, E ChIP analysis demonstrates binding of EGR1 to the promoter region of the LMTK3 gene locus in AC1 (D) and CAMA-1 (E) cells. IgG serves as a control. Error bars represent SEM of two independent experiments in triplicate; ** p< 0.01. F Dual RNA-DNA-FISH demonstrates that MIR2052HG transcripts (green signal) fail to localize to the LMTK3 gene locus (red signal) in EGR1 knockdown cells. G Hypothetical model illustrated how MIR2052HG might regulate LMTK3 transcription. The red arrows indicate the transcription direction. The transcribed MIR2052HG interacts with EGR1 protein and brings EGR1 to the LMTK3 promoter by forming DNA-RNA hydride for specific targeting of EGR1 to the LMTK3 promoter. Together with other transcription machinery, binding of EGR1 to the LMTK3 promoter initiates transcription. LMTK3 inhibits the MAPK and AKT/FOXO3 pathways, leading to regulation of ERα degradation and ESR1 transcription.

### AIs modulate LMTK3 expression in a *MIR2052HG* SNP-dependent manner

Our previous GWAS showed that *MIR2025HG* SNPs regulate its own gene expression as well as ERα expression in an estrogen or AI dependent fashion [28]. Based on our finding showing MIR2025 regulation of LMTK3, we then determined whether the expression of LMTK3 might be also *MIR2025HG* SNP- and AI-dependent using the human lymphoblastoid cell lines (LCLs) system. This cell line model system, consisting of 300 individual LCLs for which we have extensive genomic and transcriptomic data, has shown repetitively to make it possible for us to study the relationship between common genetic variant and cellular phenotypes [28,39,40]. In the presence of androstenedione, LCLs with variant genotypes for both of the *MIR2052HG* SNPs, rs4476990 and rs3802201, showed dose-dependent increases in LMTK3 expression (Fig 7A, B). However, addition of AI, either anastrozole (Fig 7A) or exemestane (Fig 7B) caused a “reversal” of the expression pattern with increased LMTK3 expression in LCLs with homozygous WT, but a marked decrease in LCLs homozygous for the variant genotypes. Of particular interest was the observation of a direct correlation between this pattern of expression for MIR2052HG and ERα [28] and that of LMTK3 (Fig 7A, B).

**Figure 7.**
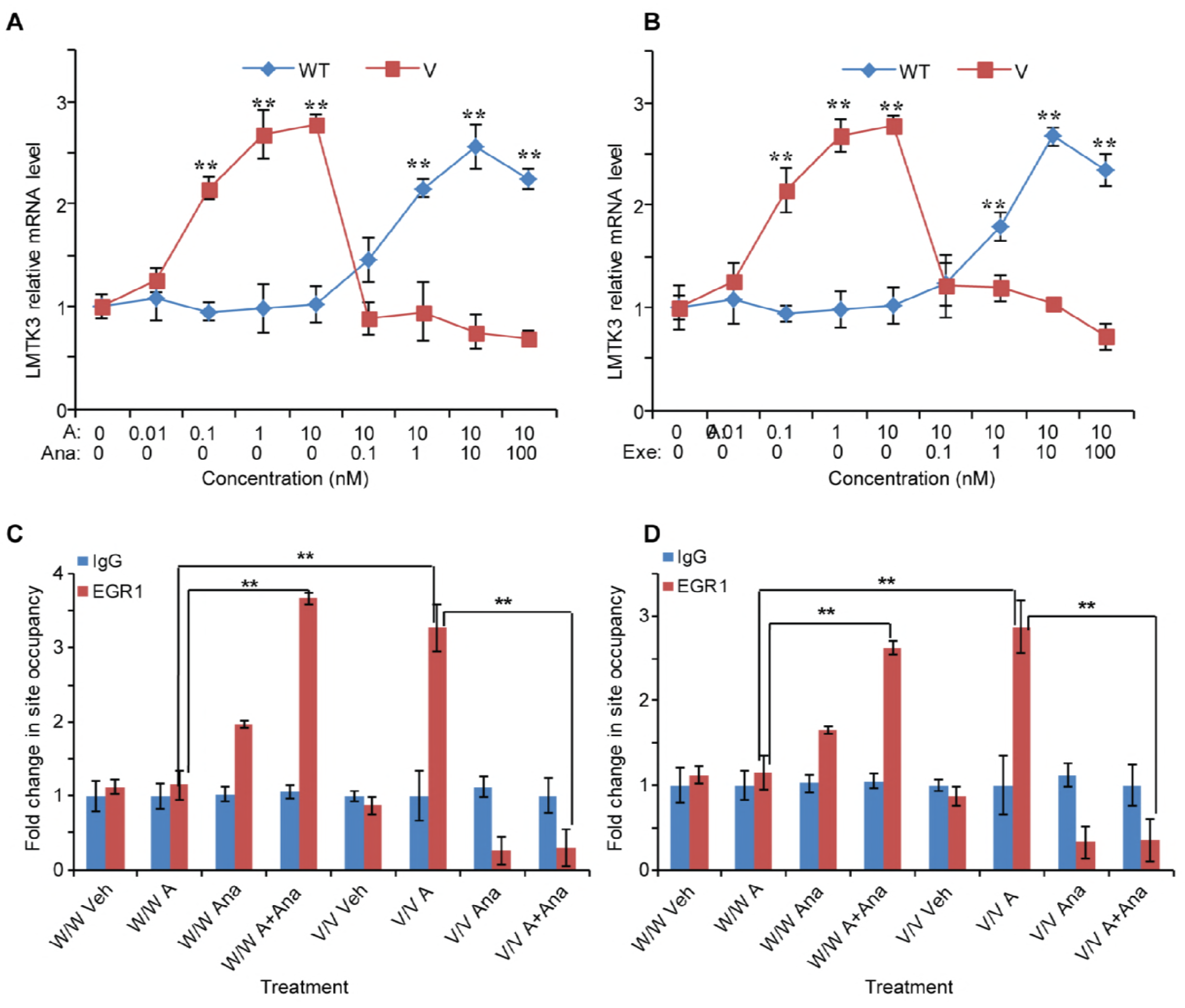
MIR2052HG-mediated SNP-dependent LMTK3 expression. A, B Androstenedione induction of MIR2052HG expression is associated with the expression of LMTK3. LMTK3 expression levels in LCLs with homozygous WT or V genotypes for MIR2052HG SNPs after exposure to androstenedione alone or with increasing concentrations of anastrozole (A) or exemestane (B). Error bars represent SEM. *P<0.05, **P<0.01. The concentrations for androstenedione (A), anastrozole (Ana), and exemestane (Exe) are indicated. C, D MIR2052HG SNPs determine androstenedione-dependent EGR1 binding to LMTK3 gene locus. ChIP assay using LCLs with known genotypes for MIR2052HG SNPs demonstrates binding of EGR1 to the promoter region of the LMTK3 gene locus after exposure to androstenedione alone or with anastrozole (C) or exemestane (D). Error bars represent SEM; ** p< 0.01. Androstenedione (A); Anastrozole (Ana); Exemestane (Exe).

Since MIR2052HG regulated LMTK3 expression in a SNP- and AI-dependent fashion (Fig 7A, B), we determined if EGR1 binding to the LMTK3 promoter region was also SNP- and AI-dependent. In the presence of androstenedione, cells homozygous for the variant SNP genotypes showed increased binding of EGR1 to the LMTK3 promoter (Fig 7C, D) relative to WT in ChIP assays using the EGR1 antibody. Anastrozole and exemestane could reverse this effect (Fig 7C, D). These results suggest that MIR2052HG facilitates EGR1 recruitment to the *LMTK3* promoter region in a SNP-dependent fashion to activate LMTK3 transcription.

## Discussions

Resistance to endocrine therapy represents a major challenge for ERα positive breast cancer therapy. Therefore, the identification of biomarkers for endocrine response and understanding mechanisms of endocrine resistance should reveal possible strategies to overcome this problem. We have previously demonstrated that germline genetic variations in *MIR2052HG* were associated with breast cancer free interval in the MA27 trial [28]. Downregulation of MIR2052HG reduced ERα positive breast cancer cell growth. The variant SNPs were associated with increased MIR2052HG expression due to increased ERα binding to EREs [28]. Therefore, MIR2052HG plays an important role in regulating ERα and endocrine resistance [28]. Recently, LMTK3, a serine-threonine-tyrosine kinase, has gained attention in breast cancer with respect to its roles in pathogenesis and therapy resistance of breast cancer [24,41,42]. The fact that overexpression of LMTK3 significantly rescued the cell growth defect caused by MIR2052HG depletion suggests that LMTK3 is one of the downstream targets of MIR2052HG (Fig 2). Furthermore, our data supported the notion that MIR2052HG *tran-*regulated LMTK3 transcription. MIR2052HG associated with EGR1 and facilitated its binding to the *LMTK3* gene promoter to activate LMTK3 expression (Figs 5 and 6), which in turn, promoted breast cancer cell growth. As a direct target of MIR2052HG, LMTK3 regulated downstream PKC/AKT/FOXO3 and PKC/MAPK/RSK1/ERα signaling, therefore, regulating breast cancer growth and AI response (Figs 3 and 4).

LncRNAs can play diverse roles in regulating gene expression as well as other cellular activities in breast cancer [43-45]. LncRNAs produce their cellular effects via several distinct mechanisms, including acting both *in cis* and *trans* [29,30]. Here, we demonstrated that MIR2052HG exerted its oncogenic role by regulating LMTK3 expression. LMTK3 is significantly elevated in high-grade breast tumors and is associated with poor survival rates in different breast cancer cohorts [24,26]. A prior study has shown that methylation is not a prevalent mechanism in the control of LMTK3 expression in breast cancer, and several somatic mutations in LMTK3 have been associated with overall survival [24]. However, we did not find any germline variations in *LMTK3* associated with breast cancer recurrence in our MA.27 cohort, suggesting a LMTK3 upstream regulator such as MIR2052HG might be the driving factor influencing this clinical phenotype. We found that MIR2052HG was induced by hormone or AIs, and it was required for the LMTK3-mediated phenotypes, including cell growth in response to AIs (Fig 7). Current research into the potential role of LMTK3 as a therapeutic target is underway [46,47]. At mechanistic level, we found that MIR2052HG positively regulated ERα at both mRNA and protein levels via LMTK3 to maintain the cancer cell growth. LMTK3 mediated the effect of MIR2052HG on AI response via ERα transcription through the LMTK3/PKC/AKT/FOXO3 signaling and protein levels via the LMTK3/PKC/MAPK pathway (Figs 3 and 4). We also found a positive correlation between the expressions of LMTK3 and ESR1 (Fig EV6) in the METABRIC and TCGA set data sample set [38], as well as in our LCLs model (p = 3.45E-04, rho=0.212). Due to the low expression levels of MIR2052HG in some of the patient samples (Fig EV3), we did not find strong correlation between the expressions of MIR2052HG and LMTK3.

EGR1 is an immediate-early gene induced by estrogen, growth factors, or stress signals [48]. The EGR1 protein binds to a specific GC-rich sequence in the promoter region of many genes to regulate the expression of these target genes including growth factors and cytokines. The mechanisms by which EGR1 activates downstream target genes appears to be cell-context dependent [49-51]. Although the DNA-binding domain of EGR1 is capable of binding to DNA through the GC-rich consensus sequence GCG (G/T) GGGCG, EGR1 can act as either an activator or a repressor of transcription through mechanisms that depend on interactions with distinct cofactors, and thus many partners, including DNA-binding proteins, have been reported to form complexes with EGR1 to activate EGR1 target gene expression [52,53]. In our study, we demonstrated that the association of MIR2052HG with EGR1 facilitated EGR1 binding to the *LMTK3* promoter (Figs 5 and 6). Based on the current data, we propose a hypothetical model that may explain how MIR2052HG contributes to LMTK3 activation and AI resistance (Fig 6G). In the model, we showed that MIR2052HG facilitated the recruitment of EGR1 to the *LMTK3* promoter through its interaction with EGR1and activated LMTK3 transcription. This process might also involve other transcription co-factors. It is possible that other proteins are also required for the binding of MIR2052HG to EGR1, since some RNA-binding proteins have been shown to be able to regulate EGR1 [54]. Nevertheless, RNA-mediated EGR1 targeting represents one mechanism by which EGR1 is recruited to its targets.

In conclusion, our findings support a model in which the protective MIR2052HG variant genotype regulates LMTK3 expression by enhancing the recruitment of ERG1 to the *LMTK3* promoter region, activating its transcription. At the mechanistic level, LMTK3 regulates ERα stability via the PKC/MEK/ERK/RSK1 axis and ERα transcription through PKC/AKT/FOXO3 pathway. This regulation may explain the effect of the MIR2052HG variant genotype on cell proliferation and response to AIs in MA.27. These findings provide new insight into the mechanism of action of MIR2052HG and suggest that LMTK3 may be a new therapeutic target in ERα positive breast cancer patients, especially those who might not respond to AIs.

## Materials and methods

### Cell lines and chemical reagents

Human ER positive breast cancer CAMA1 cell line and Human embryonic kidney cell line 293T cell line were obtained from American Type Culture Collection (ATCC). The identities of all cell lines were confirmed by the medical genome facility at Mayo Clinic Center (Rochester MN) using short tandem repeat profiling upon receipt. The breast cancer MCF7/AC1 cell line (stably transfected CYP19A1 gene) was provided by Dr. Angela Brodie (University of Maryland, Baltimore, MD). The cells were authenticated in 2015 by Genetica DNA Laboratories (Cincinnati, OH) using a StemElite ID system that uses short tandem repeat genotyping. CAMA1cells were cultured in EMEM media (Eagle’s Minimum Essential Medium) with 10% (vol/vol) fetal bovine serum (FBS) and MCF7/AC1 cells were cultured in IMEM (Improved MEM, no phenol red) with 10% heated inactive FBS in the incubator with 5% CO2 at 37°C. 293T cells were cultured in Dulbecco’s Modified Eagle’s Medium with 10% FBS in the incubator with 5% CO2 at 37°C. Five LCLs with MIR2052HG WT SNPs and five LCLs with MIR2025HG variant SNPs were selected. Before androstenedione treatment, ∼ 2×107 cells from each LCL were cultured in RPMI1640 media containing 5% charcoal stripped FBS for 24 h, followed with RPMI1640 medium without FBS for additional 24 h. Each LCL was plated into 12 well plates with RPMI 1640 medium containing 0.01, 0.1, 1 and 10 androstenedione. After 24 h treatment, increasing concentrations of anastrozole or exemestane were added. The anastrozole and exemestane concentrations ranged from 0.1, 1, 10, to 100 nM. After an additional 24 h, all LCLs were collected for further RNA extraction and qRT-PCR.

Anastrozole and exemestane (Selleckchem, Houston, TX) were dissolved in DMSO (Sigma-Aldrich) as 100 mM stock. 4-androstene-3, 17-dione (Steraloids Inc., Newport, RI) was dissolved in 100% ethanol. FLAG-ERα plasmid was provided by Thomas Spelsberg, Ph.D. (Mayo Clinic, Rochester, MN). FLAG-ERα S167A mutant was generated from the FLAG-ERα plasmid using QuickChange Site Directed Mutagenesis kit (Santa Clara, CA). HA-Ub plasmid was provided by Zhenkun Lou, Ph.D. (Mayo Clinic, Rochester, MN). LMTK3 plasmid was purchased from Origene (Rockville, MD). PKC inhibitor, Go 6983, was purchased from Sigma. U251 cells were in DMEM with 10% FBS. LCLs were cultured in RPMI 1640 containing 15% FBS. Antibodies against GAPDH, AKT, p-AKT (Ser473), FOXO3, EGR1, EZH2, RSK1, pRSK1, ERK1/2, p-ERK1/2, MEK1/2, p-MEK1/2 and Phospho-(Ser) PKC Substrate Antibody were purchased from Cell Signaling (Danvers, MA). ERα and ERα-S167 antibodies were obtained from Abcam (Cambridge, MA). LMTK3, BHLHE40, CTCF, EP300, HDAC6, POLR2A, REST, CREBBP, YY1, STAT1, SNRNP70 antibodies were purchased from Santa Cruz. CHD2 antibody was from Thermo Fisher Scientific. Secondary HRP (horseradish peroxidase)-conjugated anti-rabbit IgG and anti-mouse IgG antibodies were from Cell Signaling.

### Antisense oligo knock down and cDNA construct overexpression

Antisense oligonucleotides (ASOs) were used to knock down and study the functions of MIR2052HG. Pool of two ASOs for MIR2052HG produced with locked nucleic acids modification of 5’ and 3’ ends (Exiqon, Woburn, WA) were validated previously [28]. Negative control ASO was obtained from Exiqon (Woburn, MA). Lipofectamine RNAi Max (Invitrogen) and OPTI-MEM (Life Technologies) were used for ASOs transfection. Knock down efficiency was measured using TaqMan q-RT-PCR. The sequences of ASOs were as follows: ASO1: 5’-GTTGATTAGATTTGG-3’; ASO2: 5’-ACAGTCCCGATCTTC-3’; negative control: 5’-AACACGTCTATACGC-3’. LMTK3 plasmids (Origene, Rockville, MD) were transfected into cells using Lipofectamine 2000 reagent (Thermo Fisher Scientific, Waltham, MA). Total RNA was extracted 48 hours after transfection for RNA quantification. Whole cell lysates were collected 48 hours after transfection for western blots.

### Quantitative real-time PCR assay (qRT-PCR)

QRT-PCR assays were performed for measuring gene expression using the TaqMan RNA-to-Ct 1-Step Kit. RNA was extracted using the miRNeasy mini Kit. RNA was measured by NanoDrops300. The TaqMan primers for MIR2052HG, ESR1 and GAPDH were purchased from Life Technologies. Primers for EGR1 targeted genes were purchased from IDT. QRT-PCR reactions were prepared following manufacturer’s protocol. Samples were run using the StepOnePlus real-time PCR system.

### Western blotting

Cells were washed with cold PBS and were lysed in cold NETN buffer (100mM NaCl, 20 mM Tris·HCl pH=0.8, 0.5mM EDTA, NP-40) with Proteasome cocktail inhibitor and phosphatase inhibitor PhosSTOP EASY. Cell lysates were diluted with SDS loading buffer (SDS, glycerol, bromic acid, 1 M Tris·HCl) and boiled, centrifuged at 10,000 rpm and stored at −20 °C. Equal amounts of protein were subjected to electrophoresis in 4-15% TGX SDS gels and were transferred to PVDF membranes. Membranes were blocked in TBS with 5% BSA and 0.1% Tween-20 and then incubated overnight at 4 °C with the indicated primary antibodies. Membranes were washed with TBS-T (TBS with 0.1% Tween-20) and then incubated with HRP-conjugated anti-mouse IgG or HRP-conjugated anti-rabbit IgG for 1 hour at room temperature. All blots were visualized with the Supersignal WestPico or Supersignal WestDura chemiluminescent ECL kits and blue X-ray films or Gel Doc XR+ Gel documentation system.

### RNAseq analysis and normalization

RNA was prepared from cells using the TRIzol extraction kit. Genomic DNA was removed using the Ambion DNA-free kit. NuGEN Encore reagents were used for library preparation of total RNA samples. One microgram of total RNA input was used for each sample. The libraries were sequenced on an Illumina HiSeq 2000 sequencing system using 100-bp single-ended reads. After removing the poor-quality bases from FASTQ files for the whole transcriptome sequencing, paired-end reads were aligned by reads that were aligned to the human reference genome UCSC hg19 with Tophat 2.0.14 and the bowtie 2.2.6 aligner option. Transcript abundance was estimated using a count-based method with htseq-count.

### Cell proliferation assays

Cells were seeded (3000 cells/100 μL/well) in a 96-well plate. The CyQUANT Direct Cell Proliferation Assay kit was used to determine the cell viability in six replicates. CyQUANT assays were performed to determine the cell viability every two days. In the assay, OD 490nm (optical density) represents the absorbance at the wavelength of 490 nm. Each absorbance was normalized to the media control without any cells.

### Colony forming assays

Cells transfected with MIR2052HG ASOs or LMTK3 plasmids were plated (800∼ 1000 cells/well) in 6-well culture clusters in triplicates. Subsequently, the cells were cultured for up to 14 days at 37°C, 5% CO2 to allow colony formation. Colonies were washed with cold PBS, fixed with 4% paraformaldehyde and stained by 0.05% crystal violet. Colonies (>50 cells) were accounted with the Image J software (version 1.42q) and colony-formation rates were calculated.

### Ubiquitination assays

Approximately1.5 μg of HA-ubiquintin plasmid and 1.5 μg of pcDNA 4.1-ERα plasmid with FLAG tag were co-transfected in HEK 293T cells using Lipofectamine 2000. Twenty-four hours later, cells were reversely transfected with the MIR2052HG ASO or negative control. Approximately 2×105 ASOs transfected and control cells were subsequently seeded into each 60 mm dishes. After 64 h, MG132 was added at a final concentration of 10 μM for an additional 8 hours. Cells were then collected for the ubiquitination assay. Specifically, these cells were washed in cold PBS with NEM (1:100) and lysed in 2% SDS lysis buffer [62.5 mM Tris·HCl pH=6.8, 10% glycerol (v/v), SDS 2% (g/v)]. Immunoprecipitation assays were performed with the anti-FLAG antibody conjugated gels (ANTI-FLAG M2 Affinity gel). After washing, the FLAG gels were dissolved in 2xSDS loading buffer and boiled. These samples were then subjected to western blotting using the anti-ubiquitin antibody and anti-FLAG antibody.

### PKC kinase assay

MCF7/AC1 and CAMA-1 cells were transfected with indicated ASO or plasmids. 48 hours later, cells were lysed and the level of PKC-specific kinase activity was measured in 30 μg cell lysate using the PKC Kinase Activity Assay Kit (AbCam) as described by the manufacturer. Assays were performed in triplicates with the mean ± SD shown.

### Chromatin immunoprecipitation (ChIP)

ChIP assays were performed using EpiTect ChIP OneDay kit (Qiagen). MCF7/AC1 and CAMA-1 cells were transfected with MIR2052HG ASO for 24 hours. Cells were then subjected to CHIP assay as described by the manufacturer. LCLs were cultured in 5% charcoal stripped FBS for 24 h, followed with RPMI1640 medium without FBS for additional 24 h. LCLs were then treated with 1 nM androstenedione, 1 nM androstenedione plus 100 nM anastrozole, 1 nM androstenedione plus 100 nM exemestane for additional 24 hours. Approximately 2×10^7^ LCLs per every sample (different SNP genotypes with androstenedione or androstenedione plus anastrozole or exemestane treatment groups) were collected for the ChIP assay. Equal amount of chromatin from each sample (∼ 2 million cells each IP) and 1μg control IgG or antibody against EGR1 were used. Quantitative PCR was carried out and the result was normalized to input. All primers are listed in Table EV2.

### Fluorescent in situ hybridization (FISH)

The sequential protein staining and RNA detection were performed as previously described [55,56]. Briefly, the cells were grown in chamber slides. LMTK3 staining was performed as usual until secondary antibody is labeled in the presence of RNase inhibitor. Slides were then dehydrated by serial treatment of ethanol with different concentrations. The Alexa Fluor 488 - labeled RNA probe was obtained using the FISH Tag RNA Kit (Invitrogen). In the first step, in vitro transcription is used to enzymatically incorporate an amine-modified nucleotide into the probe template. The modified nucleotide is UTP having an NH2 group attached through a linker to the C5 position of the base. In the second step, dye labeling of the purified amine-modified RNA is achieved by incubation with amine-reactive dyes. These active ester compounds react with the primary amines incorporated into the probe template, covalently conjugating the dye to the modified nucleotide base. The purified probe is then ready for hybridization to the specimen slides at 37°C overnight. Signal was then amplified using Tyramide Signal Amplification (TSA) kit (Life Technologies). LMTK3 DNA probe was produced using the FISH Tag DNA Kit (Invitrogen). In the first step, nick translation is used to enzymatically incorporate an amine-modified nucleotide into the probe template. The modified nucleotide is dUTP having an NH2 group attached through a linker to the C5 position of the base. In the second step, dye labeling of the purified amine-modified DNA is achieved by incubation with amine-reactive dyes. These active ester compounds react with the primary amines incorporated into the probe template, covalently conjugating the dye to the modified nucleotide base. The purified probe is then ready for hybridization to the specimen. For dual RNA-DNA-FISH, we used the protocol as previously described [56]. In brief, RNA-FISH was performed by using Nick translated Alexa Fluor 488-labeled probe and followed by tyramide signal amplification kit as above. After RNA-FISH, the cells were treated by RNase A and denatured. Nick-translated BAC containing LMTK3 was labeled with Alexa Fluor 594 and used as probe. Images were obtained with the LSM 780 inverted confocal microscope runs on Zeiss’s Zen software package.

### RNA-Binding Protein Immunoprecipitation

An RNA-binding protein immunoprecipitation (RIP) assay was performed using the Magna RIP kit according to the manufacturer’s instruction. Cell lysates from 50 million cells and 2–5 μg of control IgG or antibody against BHLHE40, CHD2, CTCF, EGR1, EP300, EZH2, HDAC6, POLR2A, REST, CREBBP, YY1, and STAT1 were used. We validated the RIP assay using the SNRNP70 antibody, which can bind to U1 snRNA. Specifically, cells were washed on the plates twice with 10 mL of PBS, scraped off from plate and centrifuged at 1500 rpm for 5 minutes at 4°C and discard the supernatant. Cell pellet was re-suspended in an equal pellet volume of complete RIP Lysis Buffer, and then incubated on ice for 5 min. Dispense ∼200 μL each of the lysate into nuclease-free microcentrifuge tubes and store at −80°C. Immunoprecipitations were performed using antibodies of interest and IgG control. Anti-SNRNP70 served as controls. 50 μL of magnetic beads were washed and re-suspended in 100 μL of the RIP wash buffer. 2∼5 μg of the antibody of interest were added to each reaction and incubated with rotation for 30 minutes at room temperature. The beads were then washed three times with RIP wash buffer. 900 μL of RIP immunoprecipitation buffer were then added to each tube. The RIP lysate were thawed and centrifuged at 14,000 rpm for 10 minutes at 4°C, and 100 μL of the supernatant were added to each beads-antibody complex in RIP immunoprecipitation buffer. 10 μL of the supernatant of RIP lysate were removed as “10% input”, and stored at −80°C until starting RNA purification. The immunoprecipitations were incubated with rotating overnight at 4°C, followed by six washes with 500 μL of cold RIP wash buffer. RNA purification was then performed. Each immunoprecipitate was re-suspended in 150 μL of proteinase K buffer. The input samples were thawed and 107 μL of RIP wash buffer, 15 μL of 10% SDS, and 18 μL of proteinase K were added to the tubes and bring up the volume to 150 μL. All tubes were incubated at 55°C for 30 minutes with shaking to digest the protein, and then centrifuged briefly before being placed on the magnetic separator. The supernatant was then transferred into a new tube, together with 250 μL of RIP wash buffer. 400 μL of phenol:chloroform:isoamyl alcohol were then added to each tubes, followed by vortex for 15 seconds and centrifuge at 14000 rpm for 10 minutes to separate the phases. 350 μL of the aqueous phase were carefully removed and placed in a new tube, together with 400 μL of chloroform. After vortex for 15 seconds and centrifuge at 14000 rpm for 10 minutes, the phases were separated. 300 μL of the aqueous phase were carefully removed and place it in a new tube. 50 μL of Salt Solution I, 15 μL of Salt Solution II, 5 μL of precipitate enhancer and 850 μL of absolute ethanol were added to each tube, and kept at −80°C overnight to precipitate the RNA. The samples were then centrifuged at 14,000 rpm for 30 minutes at 4°C. The pellets were washed once with 80% ethanol, air dry, and re-suspended in 10 to 20 μL of RNase-free water. The RNAs were then analyzed by quantitative RT-PCR.

### Luciferase activity assay

Transcription activity of EGR1 was measured using the dual luciferase assay with the Cignal EGR1 Reporter Assay Kit (Qiagen). MIR2052HG knocked-down MCF7/AC1 and CAMA-1 cells were transfected with either EGR1 reporter (EGR1-responsive GFP reporter), negative control (GFP reporter construct with GFP expression controlled by a minimal promoter), or positive control (constitutively expressing GFP construct) constructs using the Lipofectamine 2000 transfection reagent. After 24 hours of transfection, luciferase assay was performed using the Dual-Glo Luciferase Reporter Assay System from Promega following the manufacturer’s protocol.

### Statistical Analysis

For cell survival, cell proliferation, kinase activity, gene expression, and quantifications, data are represented as the mean ± SEM of three independent experiments. Statistical analyses were performed with Student’s t-test. Statistical significance is represented in figures by: *P < 0.05; **P < 0.01.

## Acknowledgments

The authors acknowledge the women who participated in the MA.27 clinical trial and provided DNA and consent for its use in genetic studies. This research was supported by the Breast Cancer Research Foundation (BCRF), R01CA196648, UG1CA18967, P50CA116201 (Mayo Clinic Breast Cancer Specialized Program of Research Excellence), U1961388 (the Pharmacogenomics Research Network), the George M. Eisenberg Foundation for Charities, and the Nan Sawyer Breast Cancer Fund.

## Author contributions

J.C., J.N.I., and L.W. designed the experiments. J.C. and K.R.K. performed experiments and data analysis. M. K., L.E.S., and J.N.I. conducted MA.27 study. J.C., J.N.I., L.W., M.P.G., and R.M.W. interpreted the results and co-wrote the manuscript.

## Expanded View figure legends

**Figure EV1: LMTK3 mediates MIR2052HG -regulation of ERα.**

A Overexpression of LMTK3 in MIR2052HG knocked-down MCF7/AC1 and CAMA-1 cells did not change AKT and FOXO3 mRNA levels.

B Effects of MIR2052HG and LMTK3 on the ability of PKC to phosphorylate its substrates.

**FigureEV2: *LMTK3* DNA FISH probe map with 2 options for BACs that cover *LMTK3* gene region which were 166kb and 215kb.**

**FigureEV3: MIR2052HG and EGR1 expression in TCGA ER positive breast cancer patients.**

**FigureEV4: Knockdown of MIR2052HG specifically reduces binding of EGR1 to the *LMTK3* promoter, but not the other EGR1 targets.**

A, B Relative mRNA expression of EGR1 targeted genes after knockdown of EGR1 in MCF7/AC1 (A) and CAMA-1 (B) cells.

C, D Relative mRNA expression of EGR1 targeted genes after knockdown of MIR2052HG in MCF7/AC1 (C) and CAMA-1 (D) cells. Error bars represent SEM; ** p< 0.01, ** p< 0.05, Non-significant (NS): p> 0.05

**FigureEV5: MIR2052HG has no significant effect on other EGR1targeted genes.**

A EGR1 reporter assay in MIR2052HG knocked-down MCF7/AC1 and CAMA1 cells.

B ChIP analysis demonstrates binding of EGR1 to additional EGR1 targeted genes and knockdown of MIR2052HG has no impact on the binding. IgG serves as a control. Error bars represent SEM; Non-significant (NS): p> 0.05.

**FigureEV6: Correlations of LMTK3 expression with ESR1.**

A Correlations of LMTK3 expression with ESR1 in 2509 METABRIC breast cancer patients.

B Correlations of LMTK3 expression with ESR1 in TCGA breast cancer patients.

